# Visualizing the Costs and Benefits of Correcting P-Values for Multiple Hypothesis Testing in Omics Data

**DOI:** 10.1101/2021.09.09.459558

**Authors:** Steven R. Shuken, Margaret W. McNerney

## Abstract

The multiple hypothesis testing problem is inherent in high-throughput quantitative genomic, transcriptomic, proteomic, and other “omic” screens. The correction of *p*-values for multiple testing is a critical element of quantitative omic data analysis, yet many researchers are unfamiliar with the sensitivity costs and false discovery rate (FDR) benefits of *p*-value correction. We developed models of quantitative omic experiments, modeled the costs and benefits of *p*-value correction, and visualized the results with color-coded volcano plots. We developed an R Shiny web application for further exploration of these models which we call the Simulator of P-value Multiple Hypothesis Correction (SIMPLYCORRECT). We modeled experiments in which no analytes were truly differential between the control and test group (all null hypotheses true), all analytes were differential, or a mixture of differential and non-differential analytes were present. We corrected *p*-values using the Benjamini-Hochberg (BH), Bonferroni, and permutation FDR methods and compared the costs and benefits of each. By manipulating variables in the models, we demonstrated that increasing sample size or decreasing variability can reduce or eliminate the sensitivity cost of *p*-value correction and that permutation FDR correction can yield more hits than BH-adjusted and even unadjusted *p*-values in strongly differential data. SIMPLYCORRECT can serve as a tool in education and research to show how *p*-value adjustment and various parameters affect the results of quantitative omics experiments.

## Introduction

It is now widely appreciated there is a reproducibility crisis^1^ in biology. Although the sources of irreproducibility vary by subfield, virtually all areas of biology suffer to some degree. In quantitative transcriptomics, proteomics, or other high-throughput “omics”-type studies, a common issue is the multiple hypothesis testing problem.^2^ This problem has been known for decades and while several solutions have been put forward by statisticians, many biologists remain unfamiliar with the problem and fail to implement corrections.

In an omic screen, researchers are often looking for differences. For example, in order to find new diagnostics or therapeutic targets for Alzheimer’s disease, some studies screen for differences in protein abundances between samples from Alzheimer’s disease patients and healthy controls.^3^ In such studies, *m* analytes (e.g., genes, transcripts, or proteins) are analyzed, and the null hypothesis *H*_*0i*_ is tested against the alternative hypothesis *H*_*Ai*_ for each *i*^*th*^ analyte, where *i* ranges from 1 to *m*:

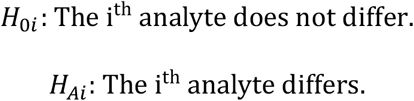

It is well known that as *m* becomes large, the probability of a false positive (rejecting a true *H*_*0i*_) becomes problematically high. Usually, *H*_*0i*_ is rejected when the *p*-value (*p*_*i*_) is below a threshold *α*, often set to 0.05; in such cases, a dataset with zero true differences (i.e., in which all *H*_*0i*_ are actually true) will typically contain *m* ∗ *α* “statistically significant” false hits nevertheless.^4^ This is the multiple hypothesis testing problem.

Several methods have been developed to address this issue and are routinely used by responsible researchers.^5,6^ However, lists of *p*-values in which correction was not applied—or for which it is unclear whether a correction was applied—are still not uncommon. In situations where uncorrected statistically significant hits are presented and are not validated with targeted experiments, such “hits” can be cited later to support false claims.

Some researchers resistant to correcting *p*-values may be concerned that a correction will decrease sensitivity—i.e., decrease the proportion of true *H*_*Ai*_ that are identified—which does occur in some multiple hypothesis corrections. Since many researchers are incentivized to maximize sensitivity, such concerns may have slowed the adoption of multiple hypothesis correction in quantitative omics.^7^ In addition, some may lack an intuitive sense of the costs and benefits of correcting omics data. We sought to address this by building an easy-to-use visual tool in the form of an R Shiny app for conceptualizing the costs and benefits of multiple hypothesis correction in situations where the true natures of the hypotheses are known. Here we describe the development of this app, which we call the Simulator of P-value Multiple Hypothesis Correction (SIMPLYCORRECT). SIMPLYCORRECT is available in the Supplementary Code and on the Internet at https://shuken.shinyapps.io/SIMPLYCORRECT/. In SIMPLYCORRECT we incorporated models designed to address the following questions:

1. What is the **benefit** of correcting for multiple hypotheses when false positives are likely?
2. What is the **cost** of data correction in datasets with many strongly differential analytes?
3. 3. What is the **dependence** of these results on the various features of this experiment, including parameters not manipulable by the scientist (e.g., effect magnitude and biological variability) and manipulable variables (e.g., number of samples and experimental variability)?
4. 4. How can the researcher **maximize** the chance of rejecting a false *H*_*0i*_ and accepting a true *H*_*0i*_ while performing responsible statistical analyses?

For these simulations we chose three correction methods common in mass spectrometry-based proteomics and transcriptomics: the Bonferroni method,^4^ the Benjamini-Hochberg procedure,^8^ and permutation-mediated false discovery rate (FDR) estimation.^9^ These methods have complementary advantages and disadvantages and cover most of the range of sensitivities and specificities of effective correction methods.^6^ Comprehensive reviews and comparisons including other correction methods can be found elsewhere.^5,6^

## Correction Methods

### Bonferroni

In the Bonferroni procedure, all *p*-values {*p*_*i*_} are simply multiplied by *m* before deciding whether to reject *H*_*0*_. (Or, equivalently, the rejection threshold *α* is divided by *m*).^4^ This reduces the family-wise error rate (FWER), the probability of obtaining at least one false hit, to the rejection threshold *α*. This method has been used successfully in trait-gene linkage screens in which a small number of genes are expected to be linked to a trait^10^ and is also appropriate in situations in which every hit will result in an action, e.g., drug approval.^11^

### Benjamini-Hochberg (BH)

For many quantitative omics methods currently employed in the study of diseases such as Alzheimer’s disease,^3^ the sensitivity of analysis using the Bonferroni method is too low to be widely useful^8^ and a higher FWER can be tolerated when hits will later be validated.^11^ As an alternative to Bonferroni, Benjamini and Hochberg developed a now classic FDR controlling method to address this. In their procedure, the *p*-values are first arranged in increasing order and multiplied by *m*/*i* where *i* is the rank of the sorted *p*-value. This product is then compared to a rejection threshold, *α*, and the following inequality is evaluated:^8^

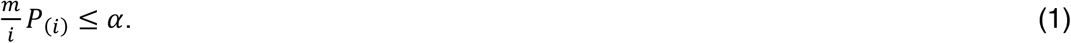

Here, the subscript *(i)* is in parentheses to reflect ascending order. The largest value of *i* for which this is true, called *k*, is determined, and then all *H*_*0i*_ for *i = 1, 2, …, k* are all rejected as long as *k* ≤ 1. If Equation (1) does not hold for any *i = 1, 2, …, m*, then no *H*_*0i*_ is rejected. Using this method, assuming accurate calculation of {*P*_*(i)*_} and mutual independence or positive regression dependency, the expected value of the false discovery proportion (FDP)—the fraction of rejected null hypotheses that are true—is equal to or less than *α*:^12,11^

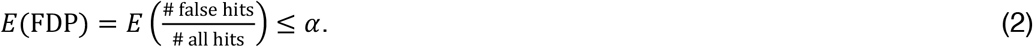

While *E*(FDP) is typically referred to by statisticians as the false discovery rate (FDR), here we refer to the FDP as FDR for accessibility and simplicity: 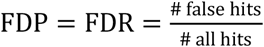. When # all hits = # all hits = 0, FDR = 0 by definition.

With BH, the *m*/*i* quotient is usually smaller in magnitude than the *m* value used by Bonferroni, allowing for the detection of more true positives while still controlling the FDR. When 0 or 1 null hypothesis is rejected, BH is equivalent to Bonferroni; when *k* is moderate or large, BH is substantially more sensitive than Bonferroni. The effectiveness, power, and ease of implementation of the BH method has made it extremely popular throughout quantitative omics communities.^13,14,15^

### Permutation

The BH procedure is unnecessarily conservative in cases where a high fraction of the null hypotheses are false, i.e., highly significant data.^11^ Rather than strict control of *E*(FDP), methods have arisen that allow a calculation of the approximate FDR of the data, allowing this approximate FDR to be lowered to a desired level (but not past it) by correcting the threshold at which *H*_*0i*_ are rejected.^11,10^ One of the most popular of these methods is permutation-mediated FDR estimation.^9^ In this method, the samples are randomly shuffled a high number of times *l* (e.g., *l* = 100). Each *j*^*th*^ time (where *j* = 1, …, *l*), statistical tests are applied (without correcting *p*-values) and the number of hits *V*_*j*_ is measured. The average number of hits across all the resulting permutations is an estimate of *V*, the number of false positives in the original permutation with samples in their proper groups.^16^ This estimate 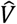 divided by the total number of hits *R* is an estimate of the FDR:^17^

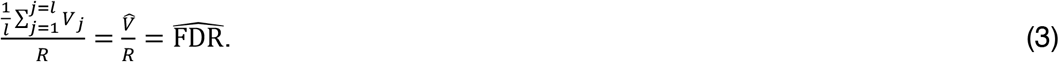

By correcting the rejection threshold, 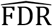 is lowered to the desired level *α*. The increased power of this method relative to BH has made it highly popular in recent years, especially in mass spectrometry-based proteomics.^14,18^

In order to compare these different methods, we sought to construct realistic models of quantitative omics experiments and use both numeric and visual metrics to assess the effects of each method on the sensitivities and specificities of the analyses.

## Results

### Simulation of P-Value Multiple Hypothesis Correction

We modeled quantitative omics experiments in R (Figure 1) and built the R Shiny app SIMPLYCORRECT which runs these omic experiment simulations with user-specified input parameters (Figure 2) (Supplementary Code). SIMPLYCORRECT simulates quantitative omics experiments in a 4-step procedure. First, analytes are generated. The user defines the number of analytes, the average true quantity of analytes in the “ome”, and the standard deviation of the distribution of analyte quantities. Here we have used Mean Quantity = 22 and Standard Deviation (S.D.) = 3 based on existing log-transformed mass spectrometry data from cerebrospinal fluid.^19^ These quantities define the “dynamic range” of the simulated ome. Analyte True Quantities are sampled from the normal distribution with µ = Mean Quantity and *σ* = S.D. Second, each analyte is “measured” in a user-defined number of samples N in the control group and test group using a user-provided Variability as a coefficient of variability (C.V) and a True Effect. This process consists of sampling the normal distribution *N*(True Quantity, [Mean * C.V.]^2^) N times to construct the control group and then *N*(True Quantity + True Effect, [Mean * C.V.]^2^) N times for the test group. For each analyte, a T-test is performed, and a *p*-value is calculated. Finally, the *p*-values or rejection threshold is corrected, and volcano plots are generated with data points color-coded according to whether each point is a false positive, false negative, true positive, or true negative. In Figure 1, True Effect = 0 was used, so that only true negatives and false positives are present.

**Figure 1.**
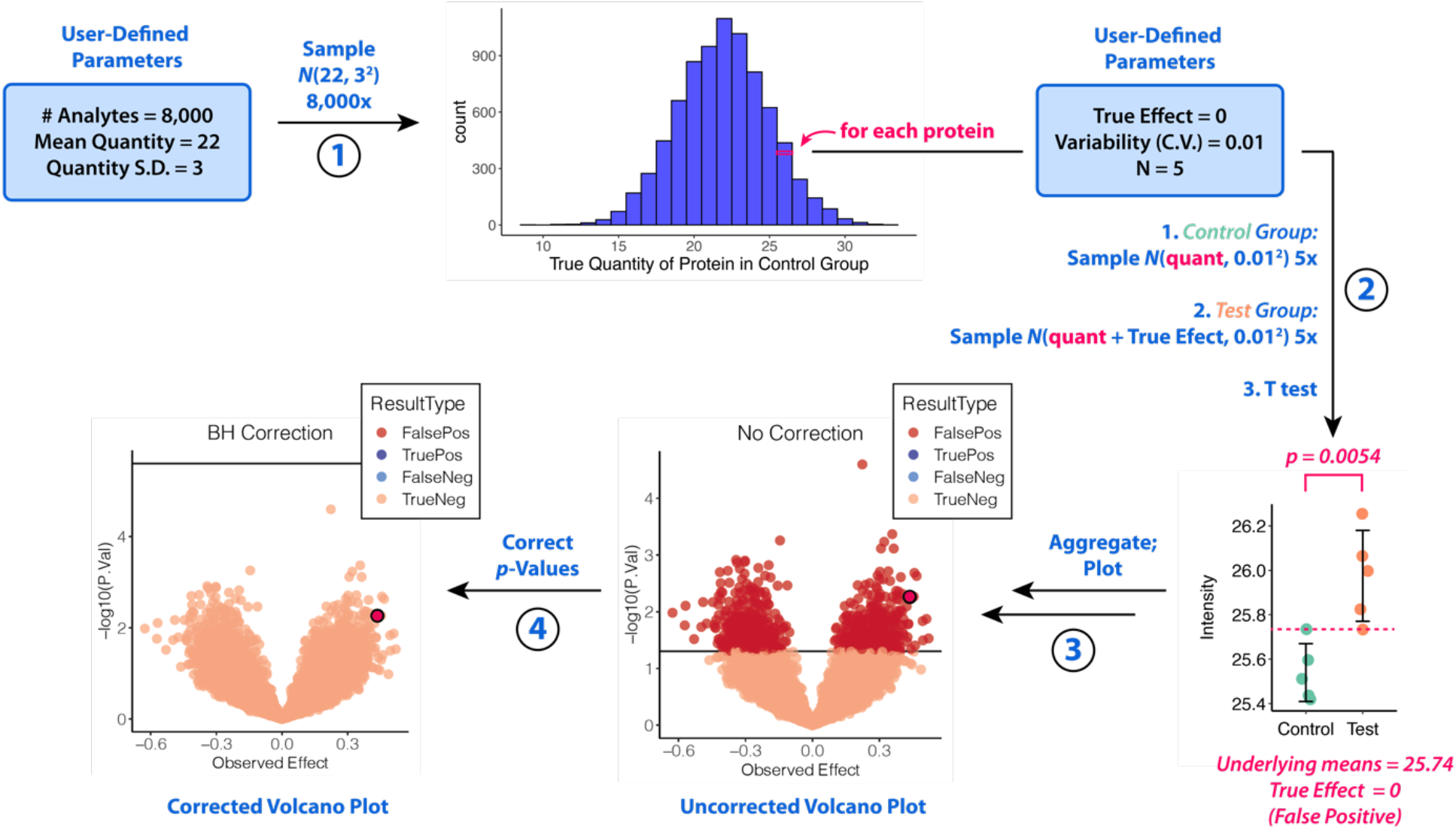
Simulator of P-Value Multiple Hypothesis Correction (SIMPLYCORRECT) for simple (non-changing or all-changing) models. Simulated quantitative omics experiments are performed in 4 steps. In this example, a True Effect of 0 was chosen, so that all nulls are true and no proteins differ between control and test groups. In Step 1, the true quantity of each protein in the control group is generated. In Step 2, each protein is “measured” N times (here 5 times) in each group with some random variability as a coefficient of variation (C.V.) (here Variability = 0.01), with the mean in the test group shifted by the True Effect (here 0). In Step 3, the results are plotted. In Step 4, the *p*-values or rejection threshold is corrected and the results are plotted. S.D. = standard deviation. *N*(µ, σ^2^) represents the normal distribution with mean μ and variance σ^2^. C.V. = coefficient of variation.

**Figure 2.**
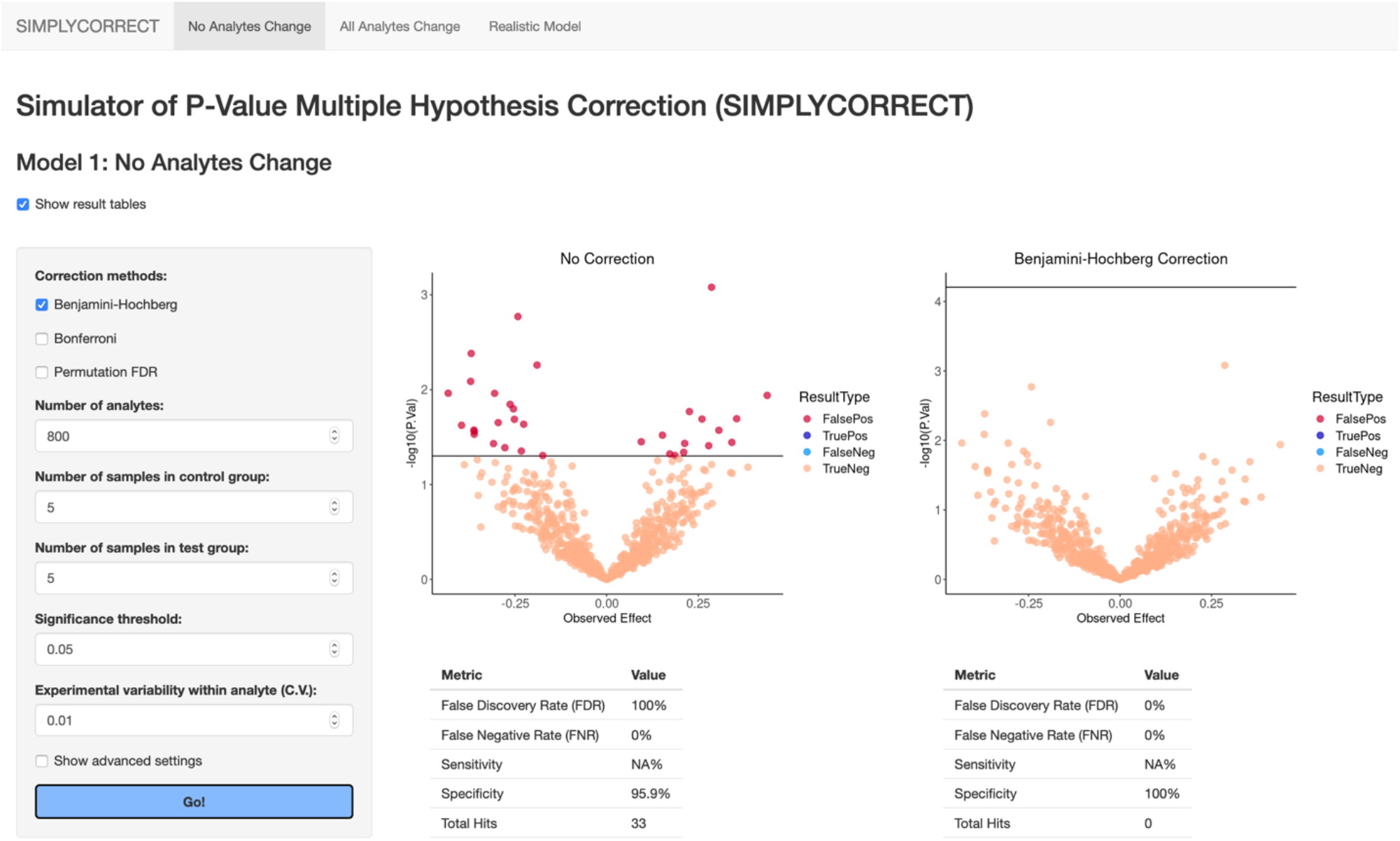
Screenshot of the SIMPLYCORRECT web application. The app is available at https://shuken.shinyapps.io/SIMPLYCORRECT/.

In the web app, the user can modify parameters and inspect how the costs and benefits of *p*-value correction vary between different experimental contexts. For this work, we ran a modified version of the SIMPLYCORRECT code to display plots from multiple different conditions side-by-side (Supplementary Code).

### Model 1: All *H*_*0i*_ Are True

First, we simulated quantitative omics experiments in which all *H*_*0i*_ are true, using values for Number of Analytes (# Analytes), Mean Quantity, Quantity S.D., and Variability (C.V.) that match log2-transformed intensity data from a real mass spectrometry-based proteomics experiment on cerebrospinal fluid (Figure 3).^19^ In this situation, there are no differences, so any effort to follow up on the significant *p*-values would be unsuccessful. While the uncorrected data in Figure 3 contain several false positives, *p*-value correction yields results that agree with the system: no *H*_*0i*_are false and none are rejected.

**Figure 3.**
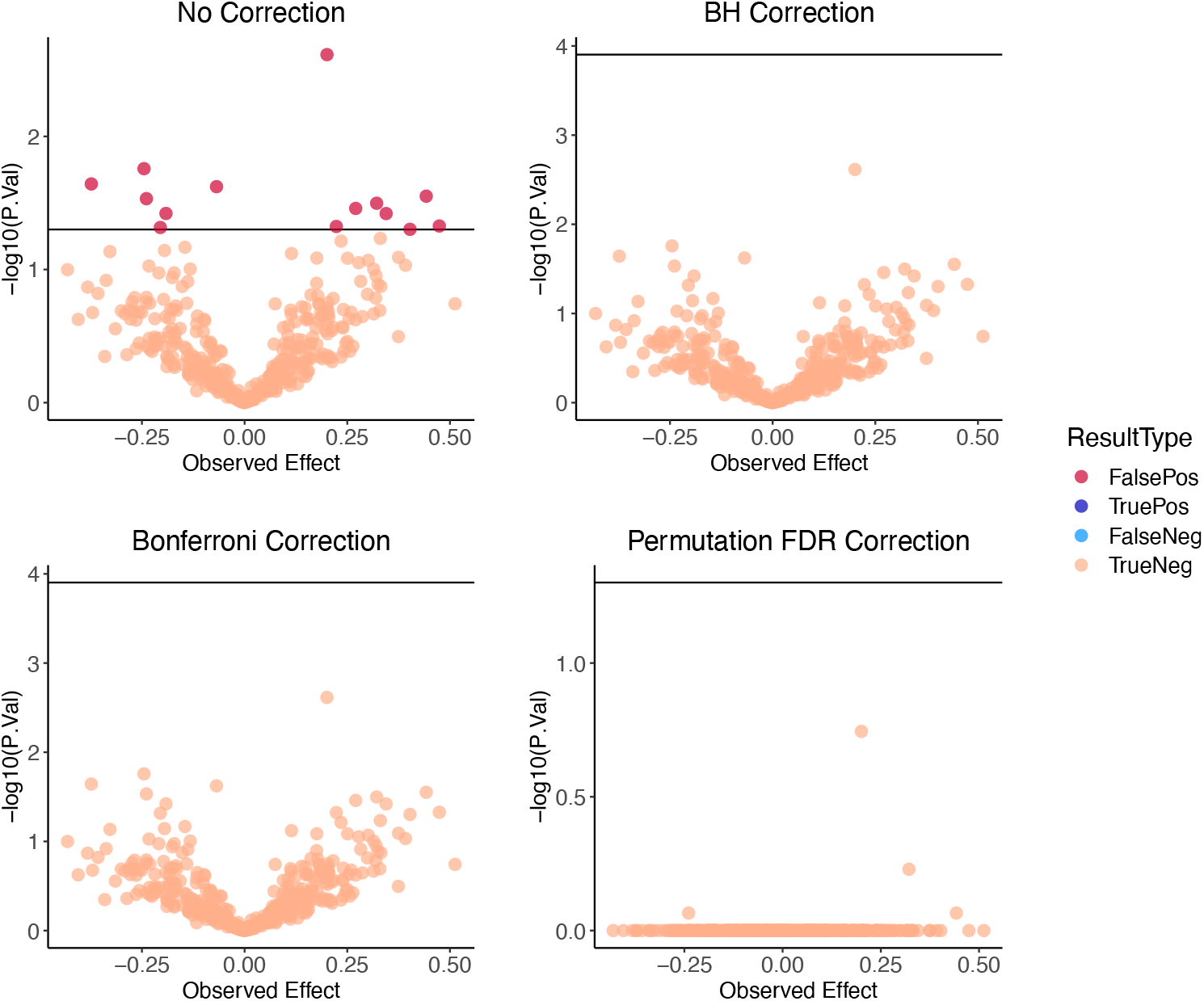
Simulated experiment in which *H*_*0i*_ is true for all genes. Parameters: # Analytes = 400, N = 3, Experimental Variability = 0.01, Significance Threshold = 0.05, True Effect = 0. BH and Bonferroni were implemented by correcting the rejection threshold while permutation FDR was implemented by correcting *p*-values. All three correction methods reduced the FDR from 100% to 0%. Note: Permutation FDR volcano plot is flattened because of cumulative minimum-based implementation in the samr package.

#### Effect of # Analytes

Increasing # Analytes (Figure 1) does not change the proportion of hypotheses that are falsely rejected without *p*-value correction (5% on average) nor the effects of *p*-value correction on the FDR in this model, but the total number of false positives increases linearly with *m*. In a 20,000-gene experiment with all true null hypotheses, for example, without *p*-value correction, the researcher would observe 1,000 false hits on average if rejecting at *α* = 0.05 or 200 false hits if rejecting at *α* = 0.01.

#### Effect of Number of Samples N

Increasing the sample size *N* also does not change the average proportion or number of hypotheses that are falsely rejected without *p*-value correction (Figure 4). In Figure 4, the BH method, as the most common method, is included exclusively for brevity. Increasing *N* does tighten the relationship between Observed Effect values and *p*-values, and observed effects get closer to the True Effect (0) as can be seen by the x-axis tick values in Figure 4.

**Figure 4.**
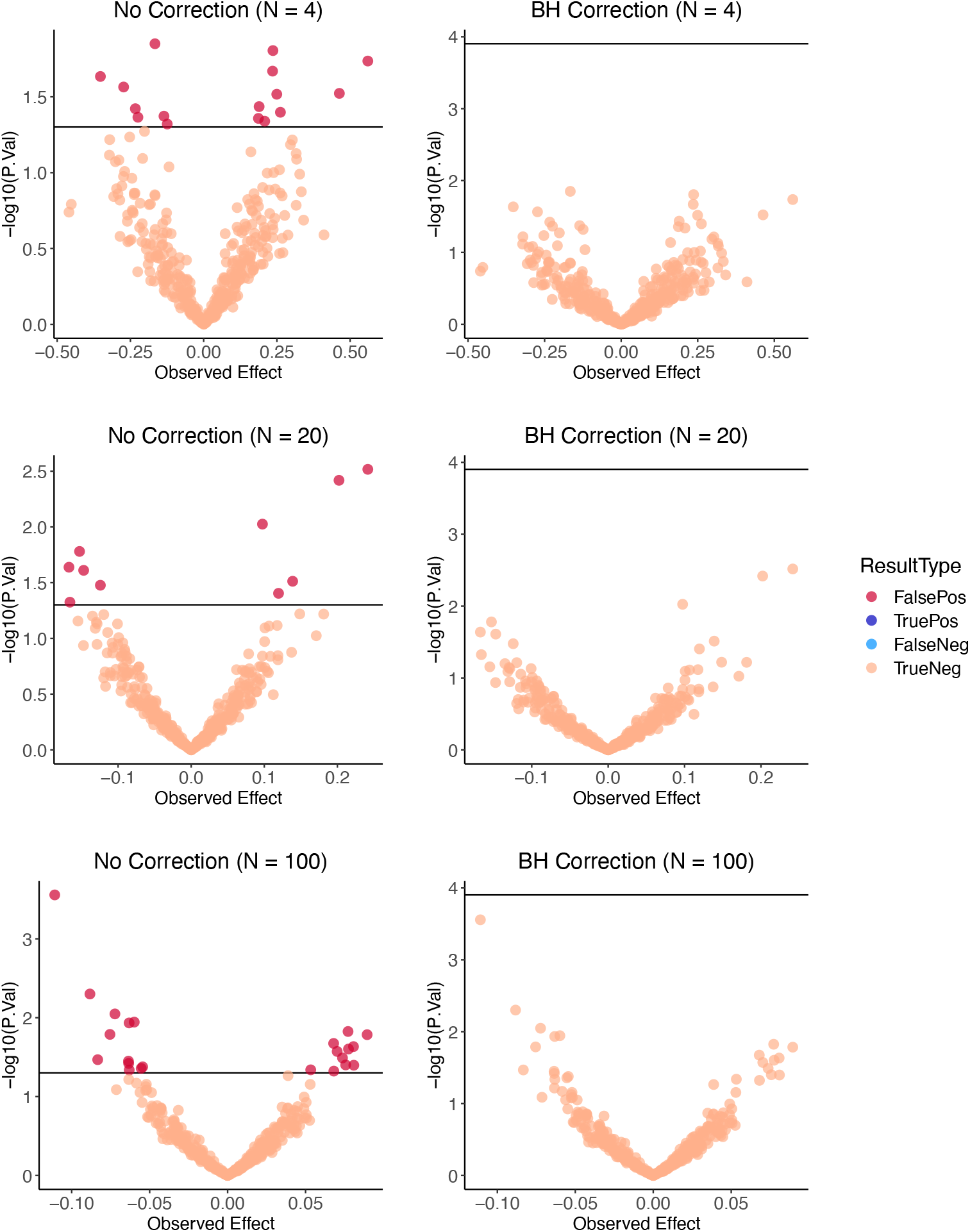
Effect of sample size N on *p*-value correction with all true *H*_*0i*_. Same parameters as Figure 3 except N. Changes in tick mark values reflect improving estimation of the True Effect (0) with increasing N.

### Model 2: All *H*_*0i*_ Are False

Next, we simulated situations in which all genes have a True Effect unequal to zero. In this model, half of the genes had some positive nonzero True Effect and the other half have the negative of that value as their True Effect. “Intensities” were “measured” incorporating normally distributed noise and the hypotheses were tested as in Model 1 (Figure 1).

#### Costs and benefits of correction methods

Figure 5 shows a typical result of a simulation with this model in which each of the 3 correction methods are applied. Though BH correction significantly reduces sensitivity in some situations, often the loss of true positive detections is minimal, as in the example shown in Figure 5 where there was only a decrease from 73% to 63%. Bonferroni, conversely, consistently results in a severe loss of sensitivity. Perhaps counterintuitively, permutation FDR correction can *increase* the sensitivity in the analysis, bringing the sensitivity to 100% in Figure 5.

**Figure 5.**
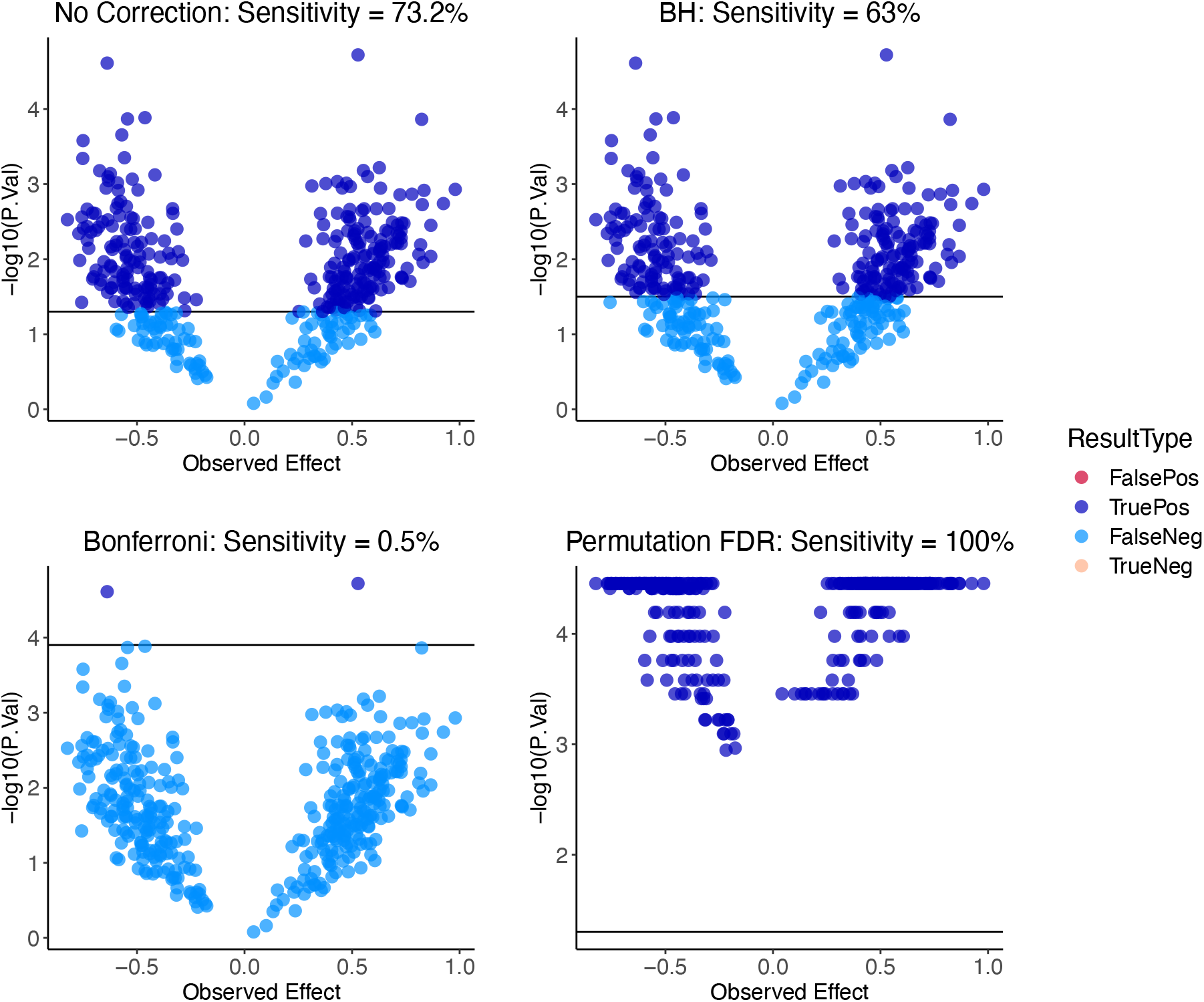
Different *p*-value corrections of data with *H*_*0i*_ all false. The following parameters were used: # Analytes = 400; True Effect = +/– 0.5; N = 4; Variability (C.V.) = 0.01; Significance Threshold = 0.05.

#### True Effect magnitude

We compared uncorrected data to BH-corrected and permutation FDR-corrected data while modifying parameters, excluding Bonferroni because of the severely low sensitivity of the method. Bonferroni can be evaluated extensively using the SIMPLYCORRECT web app.

The underlying True Effect values of the analytes had a strong effect on sensitivities (Figure 6). With weak effects, correction with BH and permutation removed most or all hits. However, with strong effects, both methods are able to detect all true positives. Importantly, *BH correction does not always create false negatives*.

**Figure 6.**
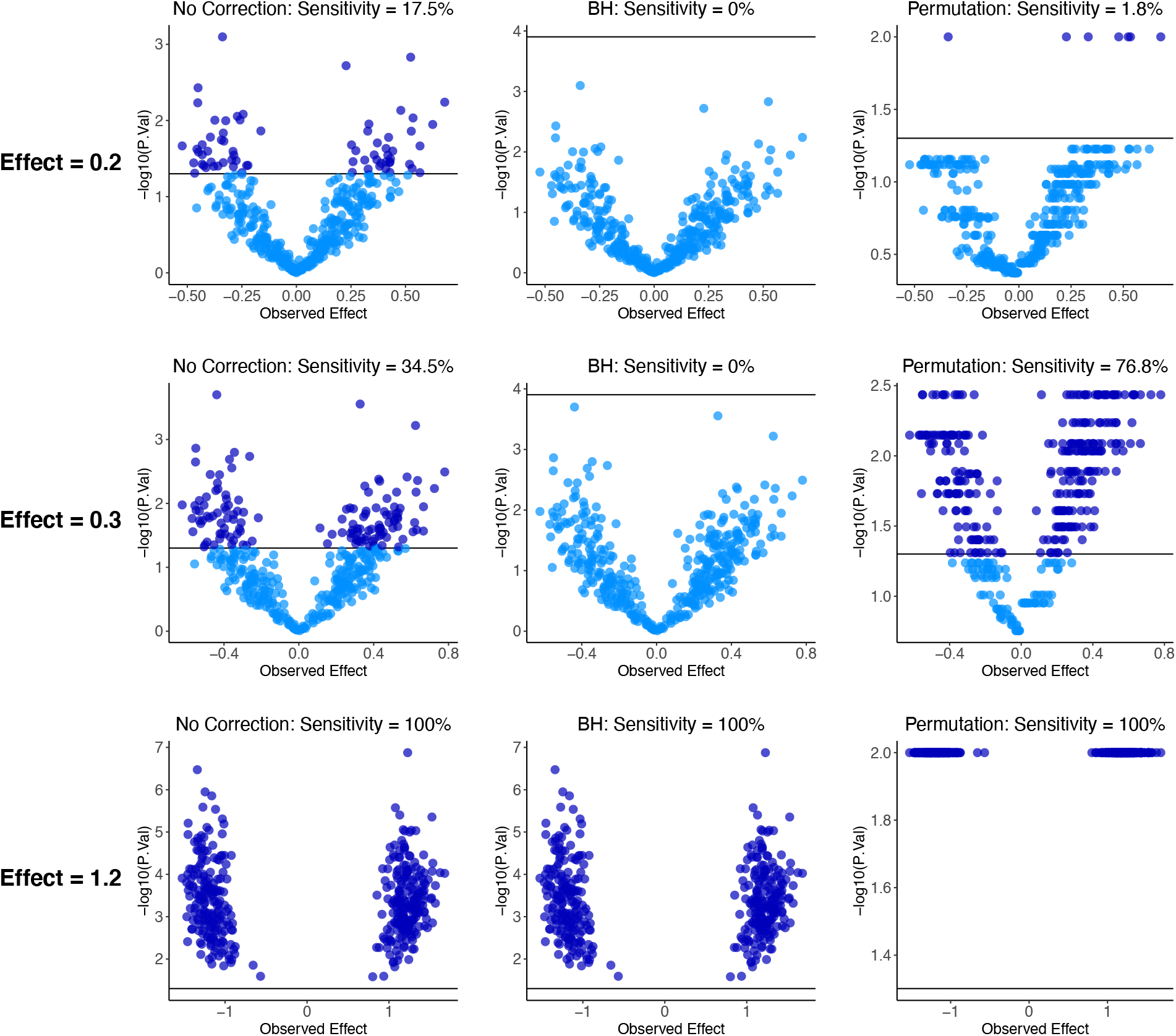
Correction of *p*-values with all false *H*_*0i*_ at various True Effects. Same parameters as Figure 5 except True Effect. Bonferroni method is excluded because of low sensitivity; Bonferroni can be examined extensively in this model using the SIMPLYCORRECT web app.

#### Effect of # Analytes

As with FDR in Model #1, the number of analytes did not affect the sensitivity in this system, though the absolute number of detected true positives increased with increasing # Analytes.

#### Effect of number of samples (N)

The number of samples (N), a parameter manipulable by the experimenter in real-life experiments, had a strong effect on the ability to detect differences while correcting *p*-values (Figure 7). Another potentially modifiable parameter, the experimental variability, had a similar effect. *With high enough N and low enough experimental variability, any false null is detectable while correcting* p*-values*, though smaller effects are harder to detect. In some contexts, increasing the effect size may also be possible, e.g., using a mouse model of Alzheimer’s disease that perturbs the system more strongly.^20^

**Figure 7.**
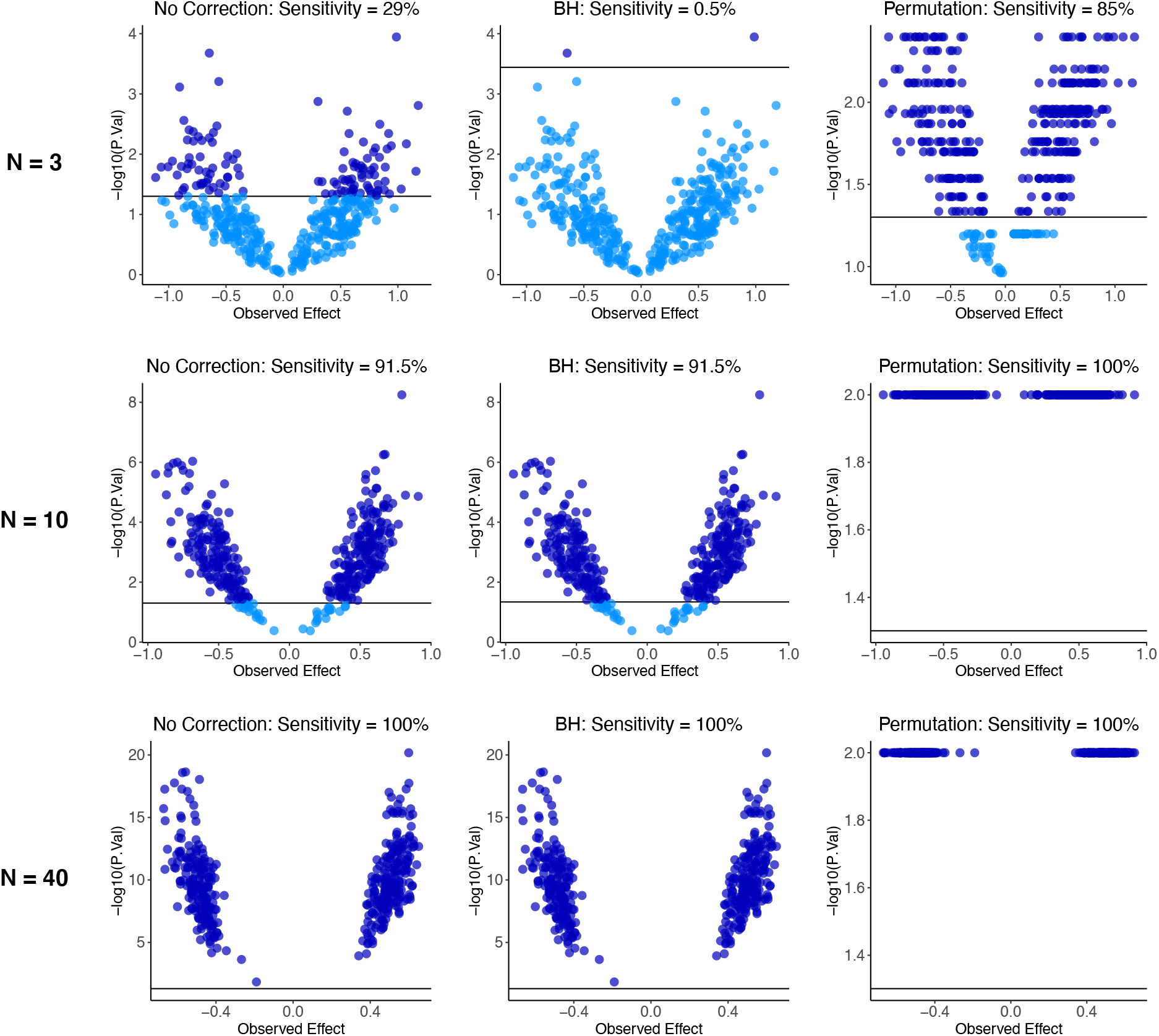
Correction of *p*-values with all false *H*_*0i*_ at various sample quantities (N). Same parameters as Figure 5 except Variability (C.V.) = 0.014 and N varies.

### Model 3: Combined True and False Nulls

#### A model of aesthetically realistic omics data

Neither of these first two models are representative of real-life data. Though more aesthetically familiar, Model 1 could be viewed as less realistic than Model 2, since perturbations rarely if ever do nothing to a biological system, and biological systems are so interconnected that perturbations may affect *all* components of a system to some small degree.

In formulating a model with both differential and non-differential analytes, early iterations overlaying Model-1-type results onto Model-2-type results appeared awkward, with two lobes of differential genes entirely separated from a clean “volcano” of non-differential genes. Implementing a normal distribution of True Effects, a more realistic alternative, raised a philosophical question: **do <u>very small</u> underlying differences, at ground truth, represent true or false null hypotheses?** We reasoned that in terms of the ultimate application of such studies, e.g., to diagnosis or treatment, they do represent the null hypothesis. In order to reconcile this with the mathematical nature of the null hypothesis, which states that nonzero effects imply *H*_*Ai*_, we found it intuitive and useful to define a “small” underlying True Effect as less than the biological variability and then set these True Effects equal to 0 (Figure 8).

**Figure 8.**
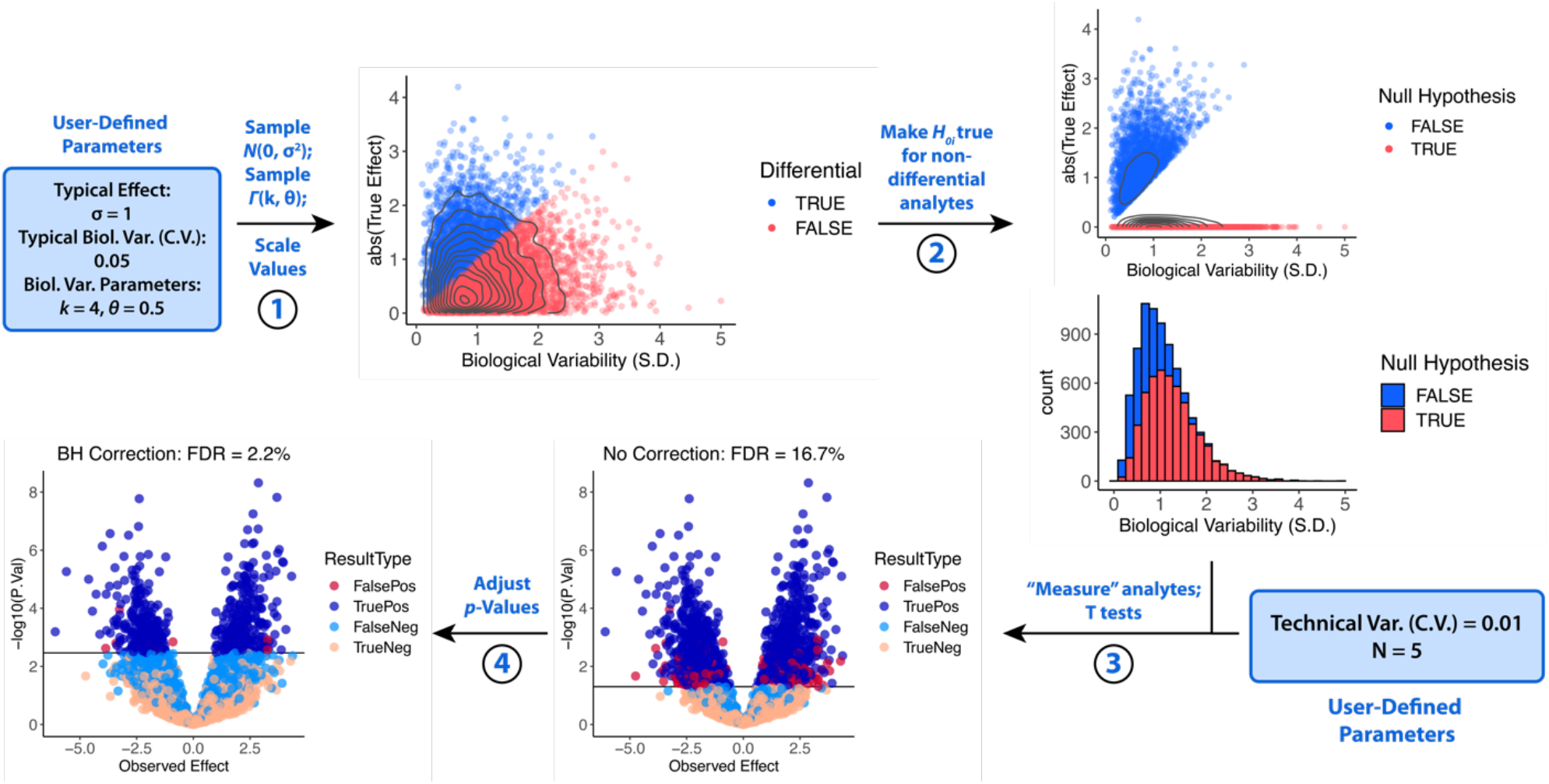
Generation of aesthetically realistic quantitative omics data by comparing effect size to biological variability. When sampling normal distributions for simulated measurements in Step 3, the standard deviation of each normal distribution is equal to the sum of biological variability and technical variability: *σ*_*measurement,i*_ = [Biol. Var. (C.V.)_*i*_ + Technical Var. (C.V.)_*i*_] * (True Quantity)_*i*_.

In Model 3, biological and technical variabilities were declared separately and then combined later into the experimental variability. Each analyte in Model 3 was assigned a random Biological Variability (C.V.) from the gamma distribution while the Technical Variability (C.V.) was constant across all analytes (Figure 8). The True Effect was then sampled from the normal distribution. If the True Effect magnitude was less than the Biological Variability S.D. (C.V. multiplied by True Quantity), *H*_*0i*_ was called true. In order to make the null hypotheses of these analytes mathematically true, the True Effects of these “non-differential” analytes were then reassigned to equal 0 before generating simulated measurements (Figure 8).

A comparison of uncorrected and corrected data in this model is shown in Figure 9. By raising the rejection threshold, BH correction reduces FDR to a low level (here 2% compared to 15.4%) while retaining several hits. Bonferroni reduces the FDR to 0%, as will happen 95% of the time (see above), at a sensitivity cost: only 2.4% of differential analytes are detected. Permutation FDR keeps the FDR at a low level (5.4% here) with significantly higher sensitivity compared to BH: here, 29.9% of differential analytes are detected.

**Figure 9.**
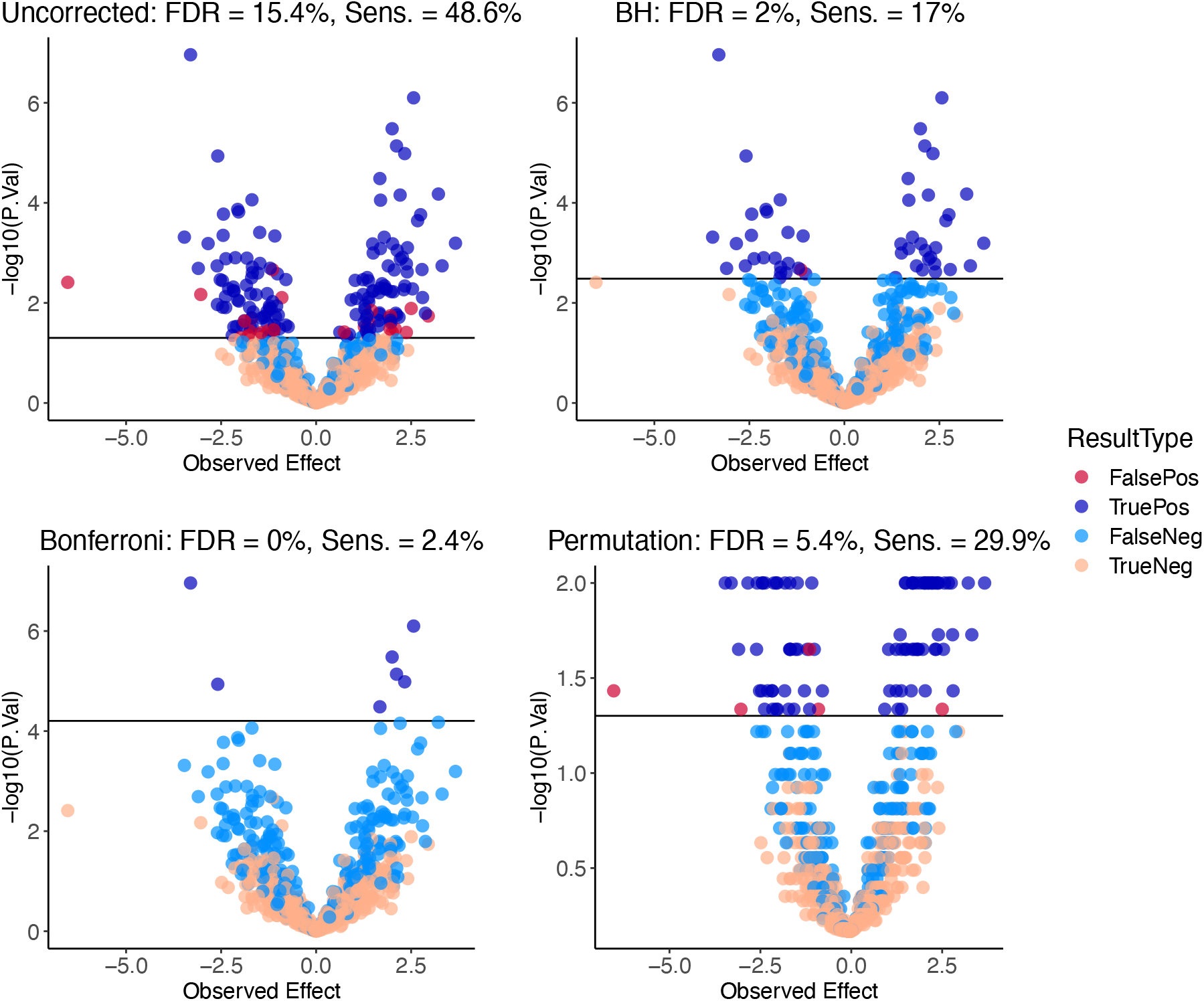
Correction of *p*-values in a realistic model. Data were generated with the following parameters: # Analytes = 800, N = 10, Typical Biol. Var. (C.V.) = 0.05, Technical Var. (C.V.) = 0.03, Typical Effect = 1, *k* = 4, *θ* = 0.5, Significance Threshold = 0.05. Sens. = sensitivity.

#### True Effect magnitudes

In order to manipulate effect magnitudes in this model, the Typical Effect was altered, changing the width of the distribution from which True Effects were sampled in Step 1 of the workflow (Figure 8). The costs and benefits of correcting *p*-values at different effect magnitudes is illustrated in Figure 10. In the case of very few changes, BH correction is necessary to lower the FDR to a safe level; occasionally, especially with high # Analytes and a poor Effect:Variability ratio, permutation can fail to control the FDR. The second example in Figure 10 is identical to that depicted in Figure 9 except with a 10-fold higher # Analytes; predictably, the FDR and sensitivity is similar in all situations, although absolute numbers are increased. With very strong effects, BH correction is less costly, and, as seen in Model 2, permutation even increases the sensitivity of the analysis relative to uncorrected data (Figure 10).

**Figure 10.**
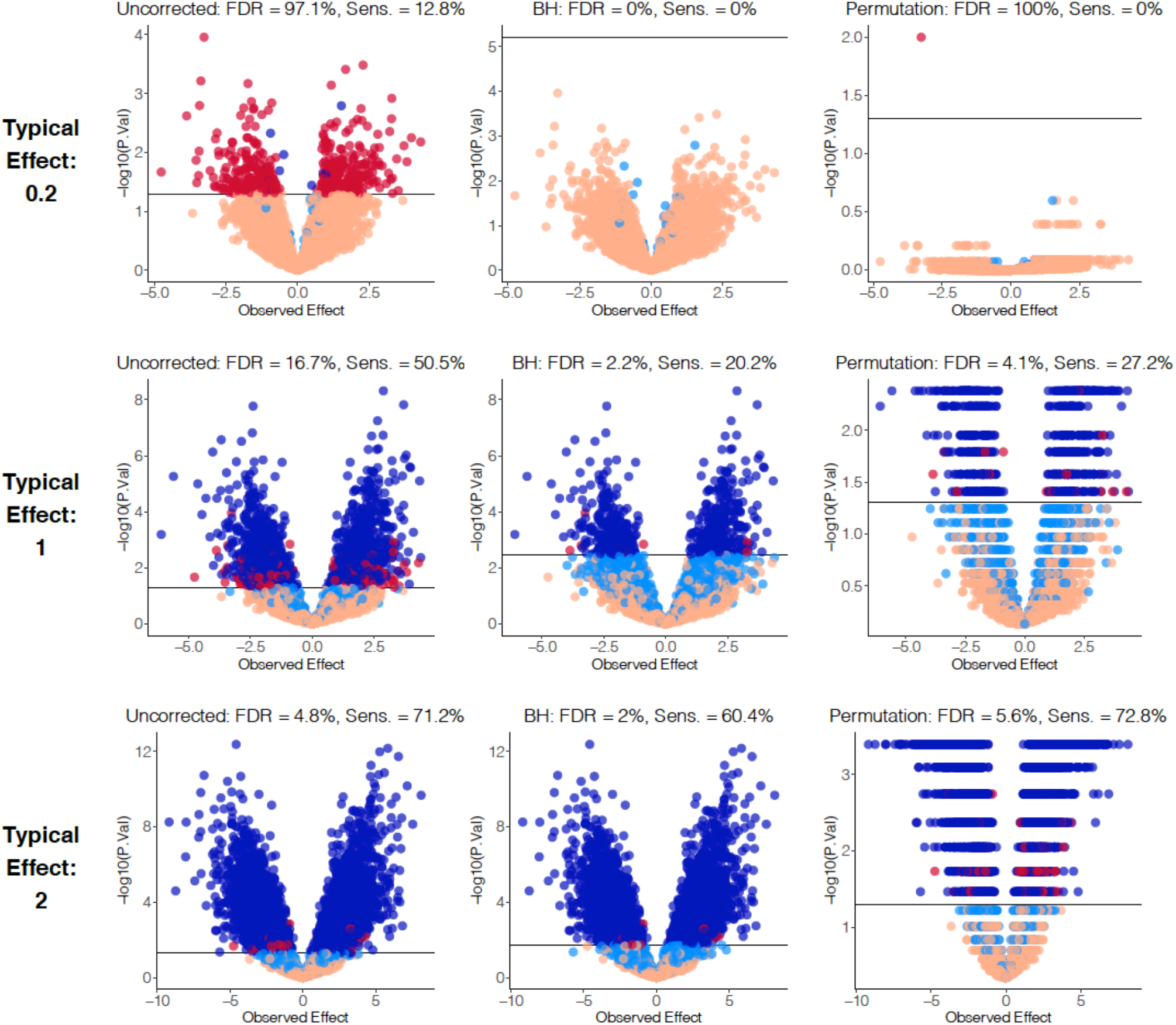
Correction of *p*-values at varying Typical Effect. Data were generated with same parameters as those used in Figure 9 except # Analytes was set to 8,000 and Typical Effects are as shown.

#### Effect of Typical Biological Variability

Lowering Typical Biological Variability values had a similar effect on the data as raising Typical Effect values, though lowering the Typical Biol. Var. had the added effect of shrinking the measured effects (x-axis in Figure 11) and tightening the relationship between measured effects and *p*-values.

**Figure 11.**
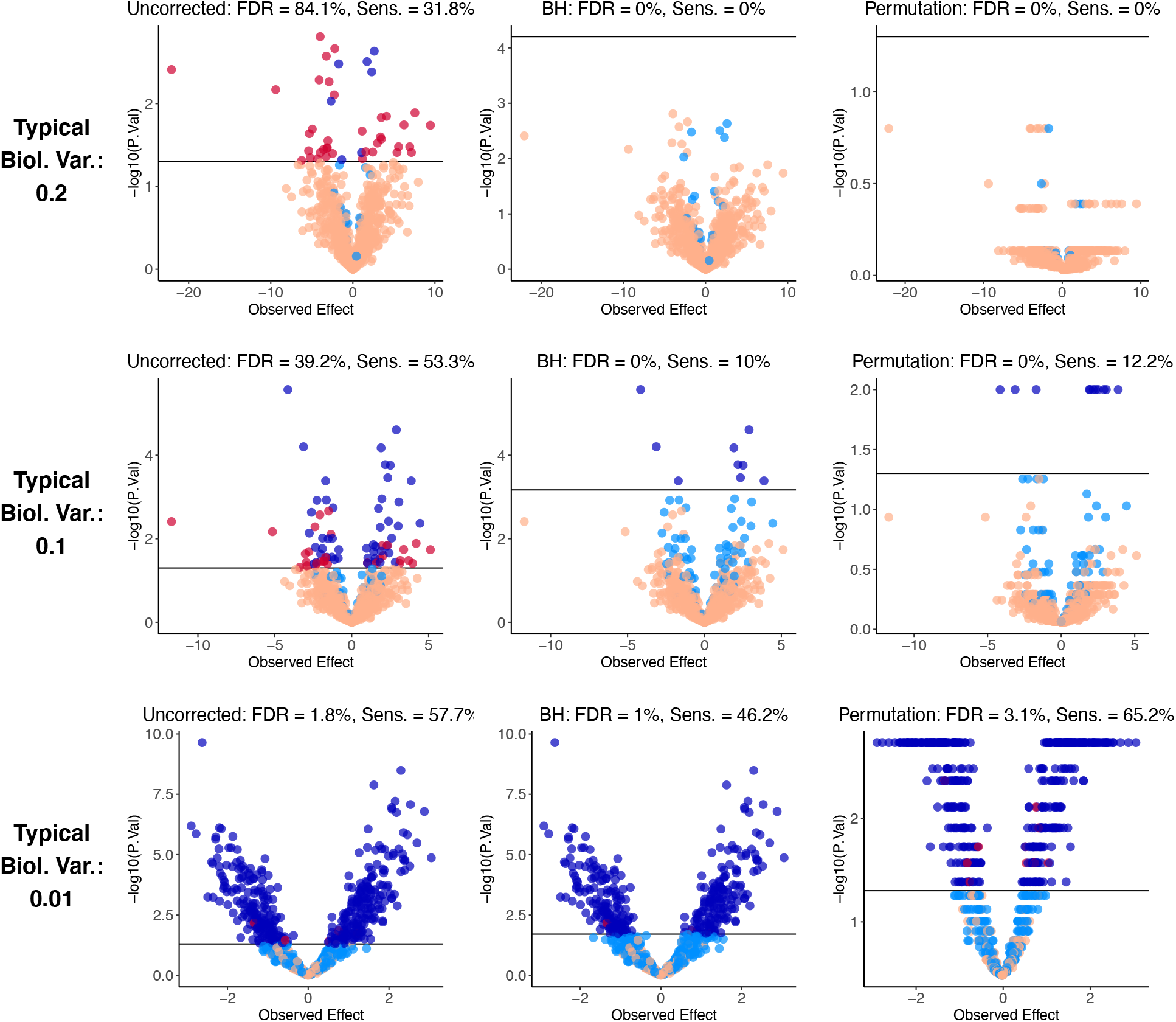
Correction of *p*-values at varying Typical Biological Variability (C.V.). Data were generated with same parameters as in Figure 9 except Typical Biol. Var.s are as shown.

#### Effect of sample number N

Parameters manipulable by the experimenter in real life were found to significantly reduce the sensitivity cost of correcting *p*-values. With high N, true and false nulls almost perfectly segregate across the corrected thresholds using both BH and permutation FDR (Figure 12). Interestingly, even at extremely high N, uncorrected can have an FDR well above the significance threshold (here, 8.4% rather than 5% at N = 100). At this extreme, corrected data, which keep the FDR at or below the threshold, can have a sensitivity higher than 95%.

**Figure 12.**
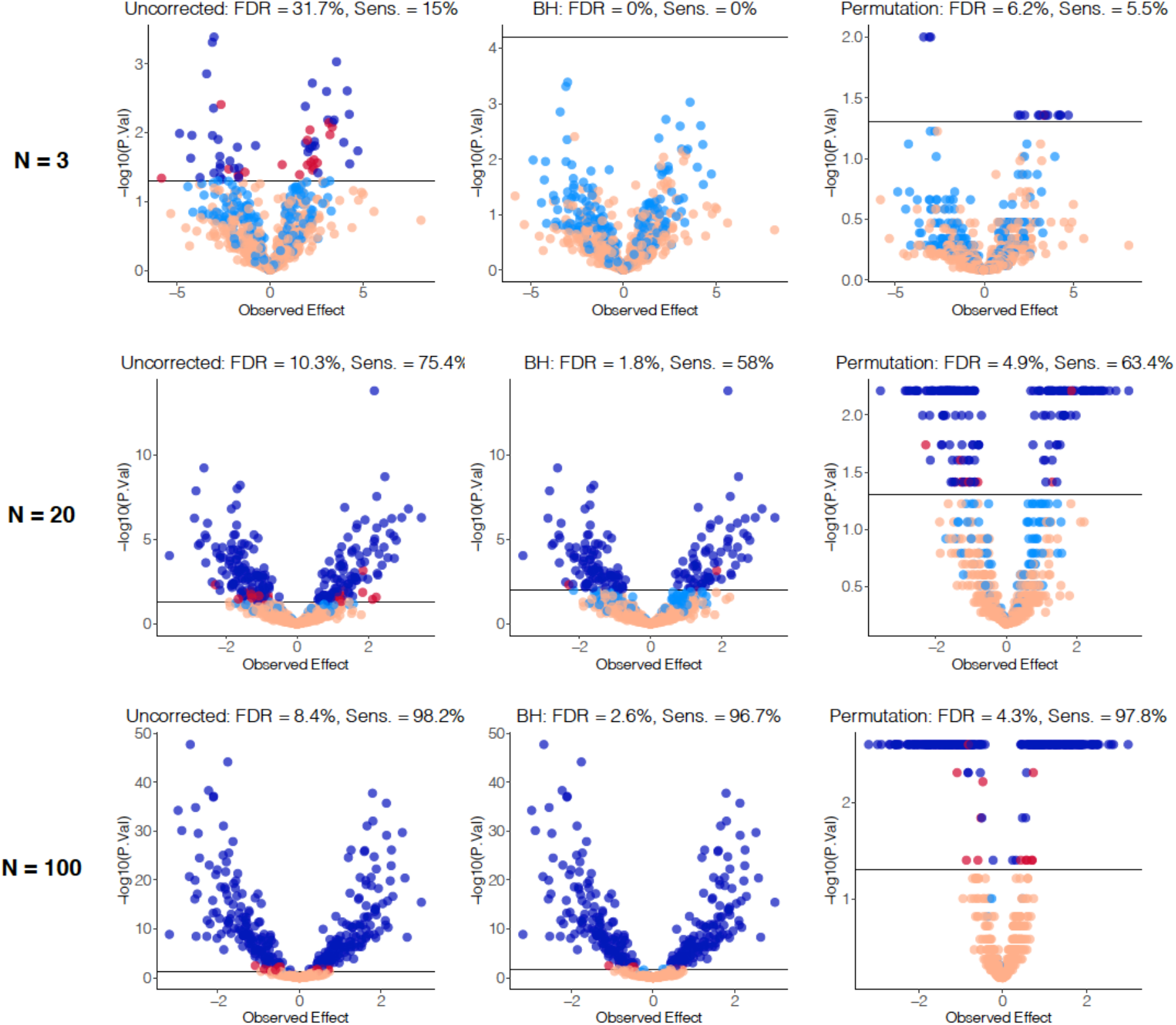
Correction of *p*-values at varying N. Data were generated with same parameters as in Figure 9 except N values are as shown.

#### Effect of Technical Variability

This model allows us to easily examine technical variability separate from biological variability. The Technical Variability parameter had a strong effect on the sensitivity and specificity of the analysis (Figure 13). In Figure 13, Typical True Effect and Biological Variability are unchanged throughout the simulations, so the numbers of true and false nulls are the same. With decreasing Technical Variability, the sensitivity of BH-corrected analysis increases from 2.7% to 51.7% to 76.5%, all with FDR below 3%. Permutation FDR had significantly higher sensitivity, especially at high Technical Variability (10.9% vs. 2.7%), with FDR ranging from 3% to 5.5%.

**Figure 13.**
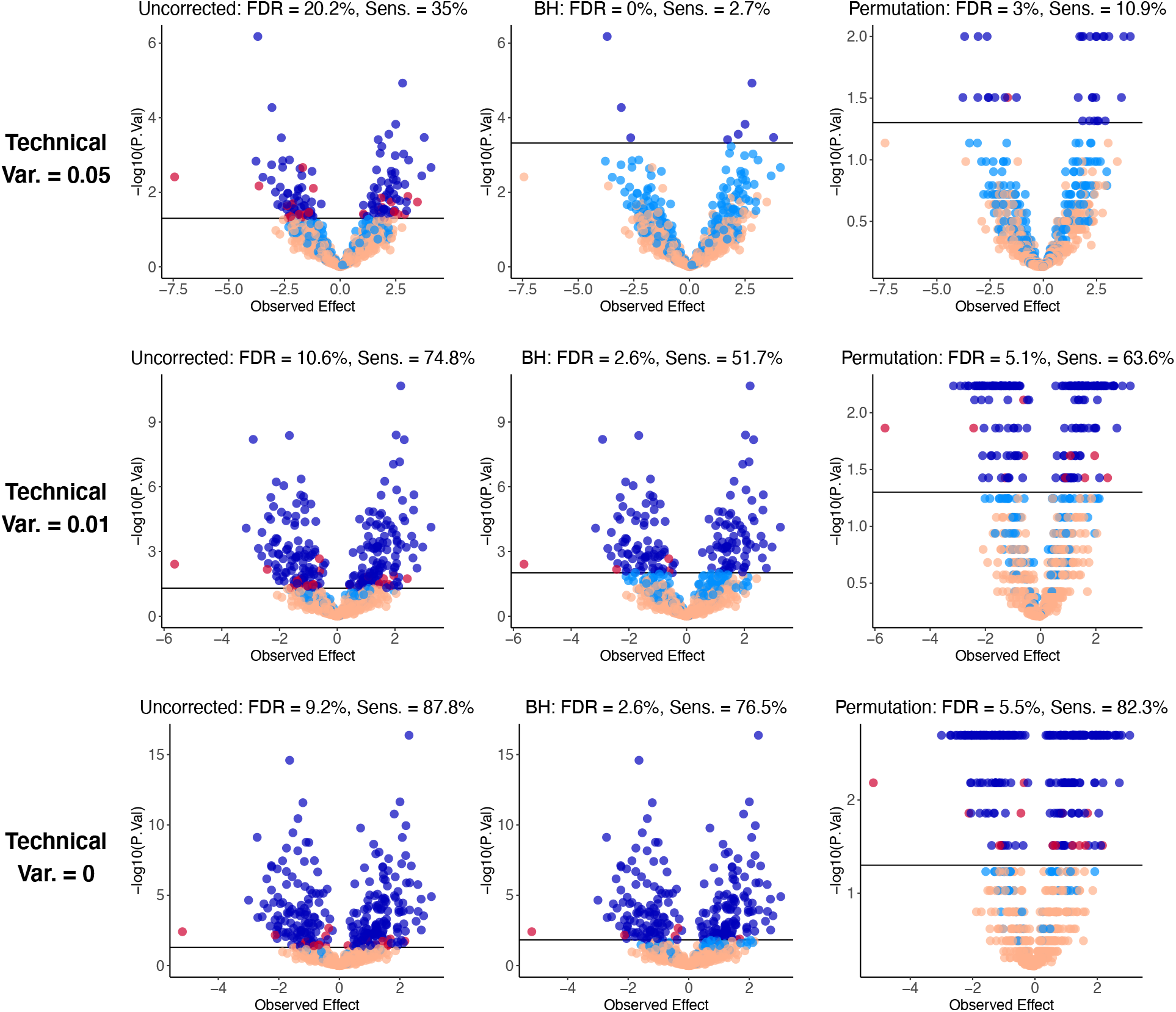
Correction of *p*-values at varying Technical Variabilities. Data were generated with same parameters as in Figure 9 except Technical Variability values are as shown.

## Discussion

To our knowledge, this is the first example of a model of quantitative omics experiments that is able to generate data that are aesthetically reminiscent of real-life data when visualized by volcano plot. In recent examples by others, True Effects were either strongly different between truly non-differential and differential analytes^5^ or the very small magnitude of technically differential True Effects sampled from continuous distributions^6^ (see above) went unaddressed; both of these approaches hampered our efforts to observe realistic results when generating volcano plots. In addition, we have built a tool that allows users to explore the parameters beyond the ranges discussed here, and we provide the computer code for researchers to adapt to their own interests and concerns (Supplementary Code).

Our models do contain several assumptions, including the mutual independence of analytes, the shapes of distributions of variables such as True Effect and Biological Variability, and the commonality of distributions (especially those shared between abundant and scarce analytes). Future work will add sophistication to our models to address these points, but in some cases, the true details are not known. Real-life distributions of biological variabilities are unknown, largely because experimental variability is a combination of biological and technical variabilities. However, conceptualizing Biological Variability as a distinct parameter is relevant to many situations, e.g., choosing a model organism when studying Alzheimer’s disease: inbred mice, outbred mice, and humans will have different biological variabilities, which should be taken into account when designing studies.^21^

In addition, increasing # Analytes in real life often means exploring additional low-abundant analytes or, e.g., in the case of mass spec proteomics, exploring additional post-translationally modified peptides or isoforms that are less likely to match a spectrum.^22^ These analytes’ distributions are unlikely to be identical to those of more common analytes; the statistical effects of deepening the exploration of the ome is still not well characterized in such scenarios. The models we present here are hoped to serve as inspiration for characterization of these ome-level features in the future.

The mutual independence of analytes may be the strongest assumption in our models, since biological omic data are often highly correlated.^19^ Dependence is known to affect the ability of correction methods to control *E*(FDP) and the FDP itself; relationships between correction procedures, error metrics, and types of dependence are still an active area of study.^23,24^ Future work will include in our simulations correlated data as well as recent correction methods, such as the usage of knockoff variables,^25^ that are designed to control *E*(FDP) in correlated data.

Despite these shortcomings, we were able to reach several reasonable conclusions regarding the questions posed in the introduction. First and foremost, it is obvious that a volcano plot with uncorrected *p*-values can look very reassuring and familiar and the hits can still be comprised entirely of false positives; the avoidance of such false hits is the main **benefit** of *p*-value correction.

In terms of **cost**, we found that BH correction does not necessarily create false negatives, and is less likely to do so with strong changes and high-quality data (low technical variability and high N). In addition, most of the time, permutation FDR is a more sensitive method than BH with minimal FDR cost. Indeed, **with very strongly differential analytes, permutation FDR can yield more hits than uncorrected *p*-values**. However, with low-quality data, permutation FDR can give a high FDR when there is a small number of hits, e.g., one hit that is false (FDR=100%), two hits with one false and one true (FDR=50%), or two false hits (FDR=100%); all of these situations were observed in our studies though only one is shown here (Figure 10). BH was reliable, with an FDR above 5% never observed (although mutual independence is a strong assumption here, as mentioned above).

Finally, in terms of the **dependence** of these results on certain parameters, most parameters affect that data in a predictable way, although it should be noted that increasing # Analytes (*m*) does not increase the quality of the data; rather, it simply increases the absolute number of all types of results while leaving the proportions unchanged. Importantly, all differential analytes (especially the most strongly changing, which are often the most biologically relevant) are detectable—while correcting *p*-values—with high enough N and low enough variability; by optimizing these parameters, the experimenter can **maximize** the sensitivity of their experiment while still correcting *p*-values.

Although we omitted the Bonferroni method from most comparisons because of its limited utility in quantitative transcriptomics and proteomics,^8^ Bonferroni, BH, and permutation FDR can all be evaluated further via the SIMPLYCORRECT app at the following URL: https://shuken.shinyapps.io/SIMPLYCORRECT/.

## Materials and Methods

The SIMPLYCORRECT R Shiny web app was coded in R using the ggplot2 and samr packages. Figures for this work were generated in R using the same packages, plus ggpubr, grid, and gridExtra. Simulated datasets were generated as described in the main text and in the SIMPLYCORRECT source code (Supplementary Code).

## Acknowledgments

The authors thank members of the Wyss-Coray Lab for helpful discussions, feedback, and support; and Dan Kluger and Art Owen for feedback and input on the manuscript. For funding and other forms of support, S.R.S. thanks Stanford BioX and the BioX Stanford Interdisciplinary Graduate Fellowship (SIGF) as well as the Stanford Center for Molecular Analysis and Design (CMAD) and the CMAD Graduate Fellowship.

